# The complex architecture and epigenomic impact of plant T-DNA insertions

**DOI:** 10.1101/282772

**Authors:** Florian Jupe, Angeline C. Rivkin, Todd P. Michael, Mark Zander, S. Timothy Motley, Justin P. Sandoval, R. Keith Slotkin, Huaming Chen, Rosa Castanon, Joseph R. Nery, Joseph R. Ecker

## Abstract

The bacterium *Agrobacterium tumefaciens* has been the workhorse in plant genome engineering. Customized replacement of native tumor-inducing (Ti) plasmid elements enabled insertion of a sequence of interest called Transfer-DNA (T-DNA) into any plant genome. Although these transfer mechanisms are well understood, detailed understanding of structure and epigenomic status of insertion events was limited by current technologies. Here we applied two single-molecule technologies and analyzed *Arabidopsis thaliana* lines from three widely used T-DNA insertion collections (SALK, SAIL and WISC). Optical maps for four randomly selected T-DNA lines revealed between one and seven insertions/rearrangements, and the length of individual insertions from 27 to 236 kilobases. *De novo* nanopore sequencing-based assemblies for two segregating lines partially resolved T-DNA structures and revealed multiple translocations and exchange of chromosome arm ends. For the current TAIR10 reference genome, nanopore contigs corrected 83% of non-centromeric misassemblies. The unprecedented contiguous nucleotide-level resolution enabled an in-depth study of the epigenome at T-DNA insertion sites. SALK_059379 line T-DNA insertions were enriched for 24nt small interfering RNAs (siRNA) and dense cytosine DNA methylation, resulting in transgene silencing via the RNA-directed DNA methylation pathway. In contrast, SAIL_232 line T-DNA insertions are predominantly targeted by 21/22nt siRNAs, with DNA methylation and silencing limited to a reporter, but not the resistance gene. Additionally, we profiled the H3K4me3, H3K27me3 and H2A.Z chromatin environments around T-DNA insertions using ChIP-seq in SALK_059379, SAIL_232 and five additional T-DNA lines. We discovered various effect s ranging from complete loss of chromatin marks to the *de novo* incorporation of H2A.Z and trimethylation of H3K4 and H3K27 around the T-DNA integration sites. This study provides new insights into the structural impact of inserting foreign fragments into plant genomes and demonstrates the utility of state-of-the-art long-range sequencing technologies to rapidly identify unanticipated genomic changes.

## Introduction

Plant genome engineering using the soil microorganism *Agrobacterium tumefaciens* has revolutionized plant science and agriculture by enabling identification and testing of gene functions and providing a mechanism to equip plants with superior traits [1, 2, 3]. Transfer DNA (T-DNA) insertional mutant projects have been conducted in important dicot and monocot models, and over 700,000 lines with gene affecting insertions have been generated in *Arabidopsis thaliana* (*Arabidopsis* henceforth) alone (reviewed in O’Malley [4]). Targeted T-DNA sequencing approaches were conducted on approximately 325,000 of these lines to identify the disruptive transgene insertions and to link genotype with phenotype. This wealth of sequence information, much of which has been made available prior to publication, is available at: http://signal.salk.edu/Source/AtTOME_Data_Source.html, has been iteratively updated since 2001, and accessed by the community over 10 million times by 2017.

The *Agrobacterium* strains used in research projects are no longer harmful to the plant because the oncogenic elements of the tumor-inducing (Ti) plasmid have been replaced by a customizable cassette that includes a diverse set of *in planta* regulatory elements. *Agrobacterium*-mediated transgene integration occurs through excision of the T-DNA strand between two imperfect terminal repeat sequences [5], the left border (LB) and right border (RB) [6], and translocation into the host genome (reviewed in Nester [7]). Hijacking the plant molecular machinery, the T-DNA is integrated at naturally occurring double strand breaks through annealing and repair at sites of microhomology [8, 9]. While the exact mechanisms behind this error prone integration are poorly understood, it is known that insertion events generally occur at multiple locations throughout the genome [5, 10]. T-DNA insertions also frequently contain the vector backbone and occur as direct or inverted repeats of the T-DNA, resulting in large intra-and inter-chromosomal rearrangements [6, 11, 12, 13, 14, 15, 16, 17, 18, 19, 20, 21]. The phenomenon of long T-DNA concatemers has previously been attributed to the replicative T-strand amplification specific to the floral dip method, and which is less often observed after tissue explant transformation of roots or leave discs [22].

Knowledge of structural variations induced by transgene insertions, including insertion site, copy number and potential backbone insertions, as well as evidence for epigenetic changes to the host genome is crucial from scientific as well as regulatory perspectives. These aspects are routinely assessed using laborious Southern blotting, Thermal Asymmetric Interlaced (TAIL) PCR, targeted short-read sequencing, or recently digital droplet PCR [4, 23]. One of the few attempts to gain deeper insight into an engineered genome was for transgenic Papaya, using a Sanger sequencing approach [14]. This work identified three insertion events, each less than 10 kilobases (kb) in length, however large repeat structures with high sequence identity are generally impossible to assemble using short-read sequences [24].

Current knowledge of structural genome changes and epigenetic stability in transgenic plants is limited. In this study we report on the genome structures of four *Arabidopsis* T-DNA floral-dip transformed plants, and for the first time we report the lengths of T-DNA insertions up to 236 kilobases along with long-molecule evidence for genome structural rearrangements including chromosomal translocations and induced epigenomic variation. To study such large insertions and rearrangements at the sequence level, we *de novo* assembled the genomes of two multi-insert lines (SALK_059379 and SAIL_232) and the reference accession Columbia-0 (Col-0) using Oxford Nanopore Technologies MinION (ONT) reads to very high contiguity. We present polished contigs that span chromosome arms and reveal the scrambled nature of T-DNA and vector backbone insertions and rearrangements in high detail. We subsequently tested transgene expression and functionality and show differential epigenetic effects of the insertions on the transgenes between the two tested vector backbones. Small interfering RNA (siRNA) species induced transgene silencing through the RNA-directed DNA methylation (RdDM) pathway of the entire T-DNA strand in the SALK-vector background, and in contrast the transgene remained active in the SAIL-line. Moreover, by profiling the occupancy of H3K4me3, H3K27me3 and H2A.Z in SALK_059379, SAIL_232 and five additional T-DNA lines, we uncovered various effects of T-DNA insertions on the adjacent chromatin landscape. In summary, new technological advances have enabled us to assemble and analyze the genomes and epigenomes of T-DNA insertion lines with unprecedented detail, revealing novel insights into the impact of these events on plant genome/epigenome integrity.

## Results

### BNG optical genome maps reveal size and structure of T-DNA insertions

To study both the location and size of T-DNA insertions into the genome, we randomly selected four transgenic *Arabidopsis* lines and assembled optical genome maps from nick-labelled high-molecular weight DNA molecules using the Bionano Genomics Irys system (BNG; San Diego, CA) from pooled leaf tissue. These Columbia-based (Col) plant lines have previously been transformed by *Agrobacterium* using different vector constructs: SALK_059379 and SALK_075892 with pROK2 (T-DNA strand 4.4 kb; [25]), SAIL_232 with pCSA110 (T-DNA strand 7.1 kb; [26, 27]), and WiscDsLox_449D11 with pDSLox (T-DNA strand 9.1 kb; [20]). The segregating plant lines resemble their Columbia parent plants showing no obvious visual T-DNA insertion-induced phenotypes. The fluorescently-labeled DNA molecules used for BNG mapping had an average length of up to 288 kb, which enables conservation of long-range information across transgene insertion sites (Table 1 and Fig. 1, 2; Supplementary Table 1). We compared assembled BNG genome maps for each of the four lines to the *Arabidopsis* TAIR10 reference genome sequence and observed up to four structural variations, due to T-DNA insertions, with sizes ranging from 27 kb to 236 kb (Fig. 1, 2, Table 1, and Supplementary Table 2). Although insertion sizes larger than the actual T-DNA cassette have been reported [6, 11, 12, 17, 20, 21], the measured length for these insertions exceeded expectations based on vector size by ∼20-60 fold. Moreover, BNG genome maps of the SAIL_232 line identified a total of seven genomic changes including three insertion events, one inversion involving ∼500 kb on chromosome 1, an inverted translocation on chromosome 3 that involves the exchange of two adjacent regions between 2.6-3.4Mb (847 kb) with 8.9-10.1 Mb (1,193 kb), as well as a swap between chromosome arm ends (chromosomes 3 and 5; Fig. 2). Previous short-read sequencing projects to identify insertion numbers (T-DNA seq; http://signal.salk.edu/cgi-bin/tdnaexpress) provided evidence for two (WiscDsLox_449D11), three (SALK_075892), four (SALK_059379) and five (SAIL_232) insertion sites, thus less than what we have observed through the BNG mapping approach (Supplementary Table S5).

**Table 1:** BNG maps and ONT assemblies identified T-DNA insertions from segregant samples. BNG maps and ONT assemblies were aligned against the TAIR10 reference genome to detect coverage, number of insertions, and maximum insertion site. Individual ONT reads identified further insertions or rearrangements that were absent from the segregant assemblies.

|  | *BNG Maps* |  |  | *ONT Assemblies* |  |  | *ONT Individual Reads* |  |
| --- | --- | --- | --- | --- | --- | --- | --- | --- |
|  | TAIR Coverage | # Insert./ Rearrange. | Max. Insert kb | TAIR Coverage | # Insert./ Rearrange. | Max. Resolved Insert kb | # Reads | # Insert./ Rearrange. |
| **SALK_059379** | 98% | 4 | 206 | 98% | 3 | 39 | 503,388 | 2 |
| **SALK_075892** | 96% | 3 | 96 | - | - | - | - | - |
| **SAIL_232** | 97% | 7 | 236 | 98% | 9 | 50 | 1,297,359 | 3 |
| **WiscDs Lox_449D11** | 94% | 1 | 164 | - | - | - | - | - |

**Fig. 1:**
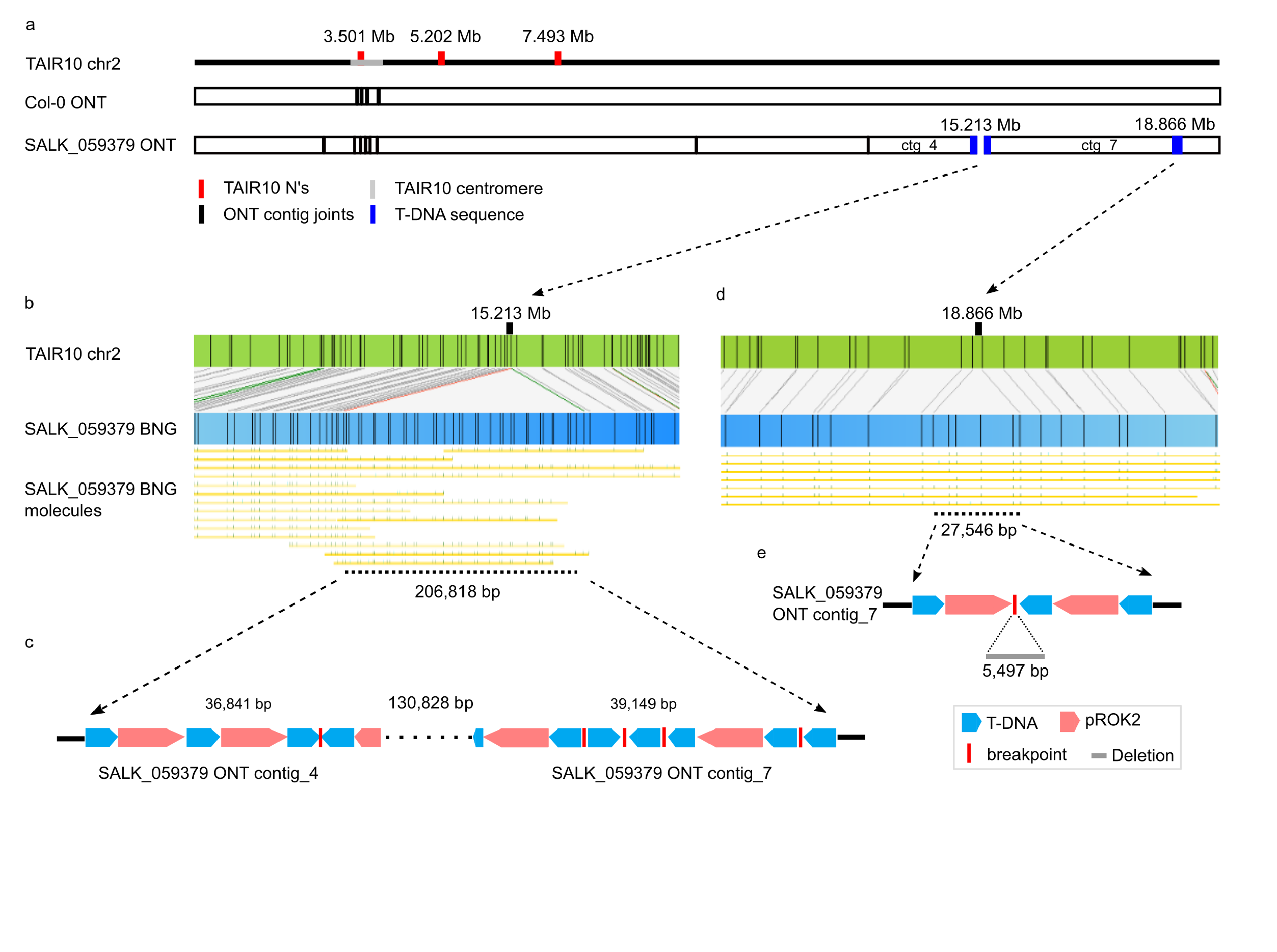
SALK_059379 T-DNA insertions are T-strand and backbone conglomerations. Schematic representation of *Arabidopsis* mutant line SALK_059379 chromosome 2 T-DNA insertions as identified from (a, c, e) ONT sequencing *de novo* genome assemblies and (b, d) BNG optical genome maps. (a) Graphical alignment of Col-0 and SALK_059379 ONT contigs to TAIR10 chromosome 2; two of the three TAIR10 misassemblies (red) are resolved within a single Col-0 ONT contig. Contig joints are represented as vertical black bars. Blue boxes indicate a broken (chr2_15Mb) and an assembled T-DNA insertion (chr2_18Mb). (b, d) represent the BNG maps that identify T-DNA insertions including an alignment of individual BNG molecules. (c) ONT contigs 4 and 7 have the chr2_15Mb insertion site partially assembled and identify a mix of direct T-DNA (blue) and pROK2 backbone (red) concatemers or scrambled stretches with breakpoints as red bars. (e) The chr2_18Mb insertion is assembled and identifies a 5,497 bp chromosomal deletion.

**Fig. 2:**
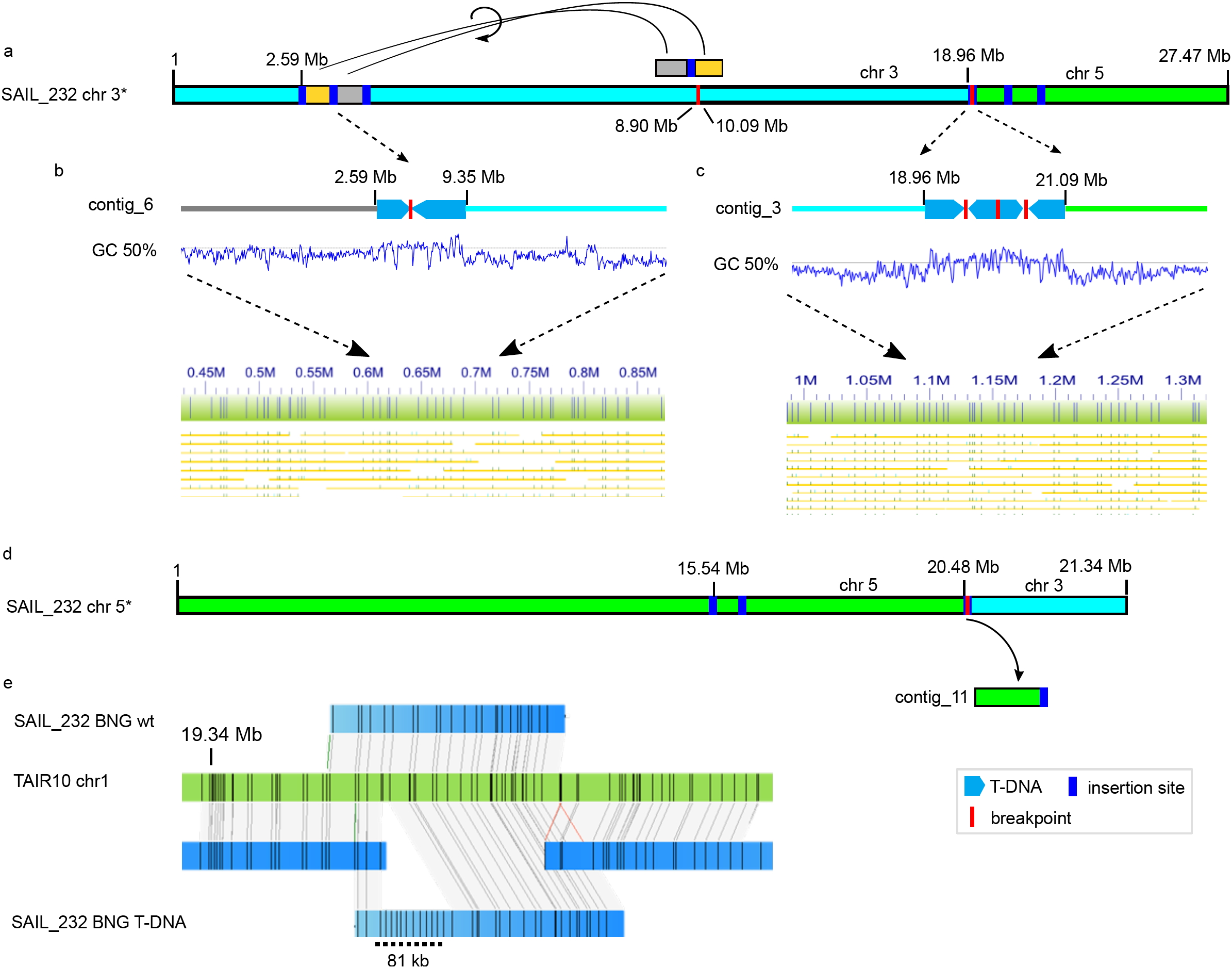
T-DNA integration induced large scale rearrangements in the SAIL_232 genome. Two T-DNA induced translocations occurred on chromosomes 3 (light blue) and 5 (lime green). (a) Inverted translocation of a 1.1 Mb segment split at a T-DNA insertion site (blue bar) was moved to a distal part of the same chromosome arm. The other chromosome arm was swapped with chromosome 5 (d). (b) The ONT assembly identified the underlying T-DNA inserts (boxed arrows) including a breakpoint (red line). (c) The chromosome swap joint site contains four pCSA110 vector fragments (boxed arrows) interspersed with breakpoints. These insertions change the CG signature in the respective region, and are confirmed by BNG alignment. (e) BNG maps (blue), aligned to *Arabidopsis* reference TAIR10 (green), are able to phase the WT (top) and T-DNA haplotypes (T-DNA; bottom) for this particular insertion site. Black lines indicate Nt.BspQI nicking sites.

### Assembly of highly contiguous genomes from ONT MinION data

The number of insertions in SALK_059379 and the type of rearrangements observed in SAIL_232 sparked our interest in analyzing these genomes at greater (nucleotide) resolution. We sequenced these engineered genomes, alongside the parent reference Col-0 plant (ABRC accession CS70000) using the Oxford Nanopore Technologies (ONT) MinION device. We performed ONT sequencing on each line using a single R9.4 flow cell (Table 1 and Supplementary Table 1) and assembled each genome using minimap/miniasm followed by three rounds of racon [28] and one round of Pilon [29]. We assembled the three lines into 40 contigs (Col-0; longest 16,115,063 bp), 59 contigs (SAIL_232; longest 16,070,966 bp) and 139 contigs (SALK_059379; longest 8,784,268 bp) (Supplementary Table 1). Individual whole genome alignments to the TAIR10 reference show over 98% coverage with 39 and 57 contigs for SAIL_232 and SALK_059379, respectively (Table 1; Supplementary Fig. 1; Supplementary Table 2).

The remaining short contigs (< 50 kb) encode only highly repetitive sequences such as ribosomal DNA and centromeric repeats that cannot be placed onto the reference. Chromosome arms are generally contained within one or two contigs, and contiguity declines with repeat content towards the centromere (Supplementary Fig. 1). Chromosome arm-spanning contigs covered telomere repeats and at least the first centromeric repeat, thus capturing 100% of the genic content. When we aligned the contigs of all assembled genomes back to the TAIR10 reference, we consistently did not cover ∼3.9Mb of the reference. The Col-0 contigs covered over 99% of the TAIR10 reference, and the only discrepancies occur at the centromeres (Supplementary Fig. 1). The high contiguity and quality of this genome assembly allowed correction of previously identified misassemblies (38/46 ‘N’-regions) in the TAIR10 reference genome (Fig. 1a, Supplementary Table 3). Our contigs were not able to span the remaining eight. BNG alignments confirmed that all ONT contigs were chimera-free, while only eleven (Col-0 and SALK_059379) or three contigs (SAIL_232) contained misassembled non-T-DNA repeats (Supplementary Table 4).

One aim of creating near complete genome assemblies was to enable the structural resolution of transgene insertions at nucleotide level, rather than with genome scaffolding alone. We next assessed contiguity at sites of T-DNA insertion. After alignment these assemblies to BNG maps we concluded that the shorter insertions, SALK:chr2_18Mb (28 kb), SAIL:chr3_9Mb (11 kb), SAIL:chr3_21Mb (25 kb; Fig. 2c) and SAIL:chr5_22Mb (11 kb) were completely assembled. Because of extensive repeats, much larger T-DNA insertions collapsed upon themselves, although contigs reaching up to 39 kb into the insertions from the flanking genomic sequences could be assembled.

### T-DNA independent chromosomal inversion in the SAIL Col-3 background

Chromosome 1 in the SAIL_232 line was assembled into a single contig (SAIL_contig_20), spanning an entire chromosome arm from telomere to the first centromeric repeat arrays (Supplementary Fig. 1). Compared with the Col-0 reference genome, we found a 512 kb inversion in the upper arm (SAIL_chr1:11,703,634-12,215,749). Because we could not find a signature of T-DNA insertion at the inversion edges, we posited that this event may have resulted from a preexisting structural variation of the particular Columbia strain used for the SAIL project (Col-3; ABRC accession CS873942). To test this hypothesis, we genotyped SAIL_232 alongside two randomly selected lines of the same collection (SAIL_59 and SAIL_107) using primers specific to the reference Col-0 CS70000 genome and the SAIL-inverted state (see Methods). PCR analysis confirmed that this “inversion” was common to all three independent SAIL-lines tested, and absent from Col-0 CS70000 (Supplementary Fig. 2). Thus the event was not due to the T-DNA mutagenesis, rather is an example of the genomic “drift” occurring during the propagation Columbia “reference” accession within individual laboratories.

### SALK_059379 T-DNA insertions are conglomerates of T-strand and vector backbone

To annotate T-DNA insertion sites within the assembled genomes, we searched for pROK2 and pCSA110 plasmid vector sequence fragments within the assembled contigs (Supplementary Table 5). The assembled SALK_059379 genome contained three of the four BNG map identified T-DNA insertions: SALK:chr1_5Mb, chr2_15Mb and chr2_18Mb (Table 1, Fig. 1 a,c,e). Specifically, SALK:chr2_18Mb, the shortest identified insertion with 28,356 bp, was completely assembled (contig_7:3,690,254-3,719,373) and included a genomic deletion of 5,497 bp (chr2:18,864,678-18,870,175). Annotation of the insertion revealed two independent insertions; a T-DNA/backbone-concatemer (11,838 bp) from the centromere proximal end, and a T-DNA/backbone/T-DNA-concatemer (16,463 bp) from the centromere distal end, both linked by a guanine-rich (26 G bases) segment of 55 bp (Fig. 1e). Two independent insertion events, 5,497 bp apart, potentially created a double-hairpin through sequence homology, that was eventually excised and removed the intermediate chromosomal stretch (Supplementary Fig. 3).

The second and third insertions, SALK:chr1_5Mb (131 kb) and SALK:chr2_15Mb (207 kb), were partially assembled into contigs. SALK:chr1_5Mb contig_10 and contig_5 contain 25,132 bp and 33,736 bp T-DNA segments. Similarly, the extremely long chr2_15Mb insertion was partially contained within contig_4 and contig_7 (Fig. 1c), leaving an unassembled gap of 131 kb. The recovered structure of this insertion is noteworthy as it represents a conglomerate of intact T-DNA/backbone concatemers, as well as various breakpoints that introduced partial vector fragments with frequent changes of the insertion direction (Fig. 1c). Finally, the fourth insertion SALK_059379 (SALK:chr4_10Mb) was absent from the assembly. However, we observed a single ONT read (length 10,118 bp) supporting the presence of an insertion at this location. We also recovered a further ONT read (length 15,758 bp) that anchors at position chr3:20,141,394 and extends 14,103 bp into a previously unidentified T-DNA insertion (Supplementary Fig. 4). PCR amplification from DNA samples using the genomic/T-DNA junctions sequences from segregant and homozygous seeds, in fact, confirmed the presence of all five insertions, revealing heterozygosity within the ABRC-sourced seed material.

While BNG maps were successful in placing long “T-DNA only” sequence contigs into the large gaps (e.g. SALK:chr1_5Mb), the four “short” T-DNA-only sequence contigs of ∼50 kb or less did not contain sufficiently unique nicking pattern to confidently facilitate contig placement.

### Large-scale rearrangements reshape the SAIL_232 genome

We next searched the SAIL_232 ONT contigs for pCSA110 vector fragments, and were able to confirm all BNG-observed genome insertions (Table 1). This search additionally identified a further T-DNA insertion at chr5:20,476,509 (Supplementary Table 5) that was not assembled in the BNG maps.

We found that chromosome 3 harbored two major rearrangements (Fig. 2). The first was a translocation of a 1.19 Mb fragment (chr3:8,902,305-10,095,395), which split at an internal T-DNA insertion at reference position 9,343,053 bp (Fig. 2a). The resulting two fragments were independently inverted prior to integration just before position chr3:2,586,494 (Fig. 2 a,b). The second major change was a swap between the distal arms of chromosomes 3 and 5, which is supported by two SAIL_232 BNG genome maps as well as two ONT contigs (Fig. 2a-d). Here, chromosome 3 broke at 21,094,402 nt, and chromosome 5 at 18,959,379 nt and 20,476,664 nt, and the larger chromosomal fragments swapped places. Specifically, this translocation was captured in SAIL_contig_31, showing the fusion of chr5:20,476,664-end to chromosome 3 after position 21,094,402 nt. The reciprocal event joined (SAIL_contig_2) chromosome 3 fragment (21,094,407-end), almost seamlessly to chromosome 5 after reference position 18,959,379 nt (Fig. 2c,d). The genomic location of an excised fragment of chromosome 5 (chr5:18,959,380 - 20,476,663 found within SAIL_contig_11), was not determined (Fig. 2d).

Finally, the 81-kb insertion at SAIL:chr1_19Mb was identified from a BNG contig that aligned together with a contig that did not harbor the insertion (Fig. 2e). This apparent phasing of this heterozygous region was the only occasion where we observed this. The insertion consists of four tandem T-DNA copies (∼30 kb) followed by ∼20 kb of breakpoint interspersed T-DNA and vector backbone was partially assembled (50,676 bp) at the 5’ end of contig_5 (Fig. 2e). Although not assembled as part of the flanking SAIL_contig_47, we recovered single ONT read that contain T-DNA as well as genomic DNA sequences. In summary, the BNG maps perfectly aligned with the ONT assembly of the T-DNA insertion haplotype (Fig. 2e).

### T-DNA Integration occurs independently from both double-strand break ends

While both sequenced lines share similar numbers of T-DNA insertion events, the genome of the SAIL_232 plant line underwent more significant changes to its architecture. All genome insertion sites began and ended with the LB of the T-DNA strand, providing evidence for independent transgene integration at both ends of the DNA double-strand break. We did not recover any LB sequences at the chromosome/T-DNA junction, in line with literature reports that usually 73 - 113 bp are missing from the LB sequence inwards [15, 30]. Internal T-DNA sequence deletions were also seen at breakpoints within the insertion (Fig. 1c, Fig. 2c). As observed for the SALK:chr2_18Mb chromosomal deletion (Fig. 1e and Supplementary Fig. 3), we cannot exclude that long homologous stretches between the independently inserted T-DNA/vector backbone concatemers represent inverted repeats.

### Transgenes are functional in SAIL-lines but are silenced in SALK-lines

We next wanted to assess the effects of T-DNA insertions on the epigenomic landscape. The pROK2 T-DNA strand contains the kanamycin antibiotic-resistance gene *nptII* under control of the bacterial nopaline synthase promoter (NOSp) and terminator (NOSt), and an empty multiple cloning site under control of the widely used Cauliflower mosaic virus 35S (CaMV 35S) promoter and NOSt. The CaMV 35S constitutive overexpression promoter has previously been described to cause homology-dependent transcriptional gene silencing (TGS) in crosses with other mutant plants that already contain a CaMV 35S promoter driven transgene [31, 32]. In germination assays, we confirmed that the kanamycin selective marker is not functional in SALK lines propagated for more than a few generations [4]. While ∼75% of SALK_059379 seed initially germinated on kanamycin-containing plates, these seedlings stopped growth after primary root and cotyledons emergeance (Fig. 3a). The SAIL_232 pCSA110 T-DNA segment encodes the herbicide resistance gene *bar* (phosphinothricin acetyltransferase), under control of a mannopine synthase promoter [26]. In contrast to SALK_059379, we confirmed proper transgene function by applying herbicide to soil-germinated plants [33, 34] (Fig. 3b). We corroborated these differential phenotypes by mapping RNA-seq reads to the corresponding transformation plasmid and found that the SAIL_232 *bar* gene was expressed, while in SALK_059379 the *nptII* gene was not expressed, most likely due to epigenetic silencing (Fig. 3c,d).

**Fig. 3:**
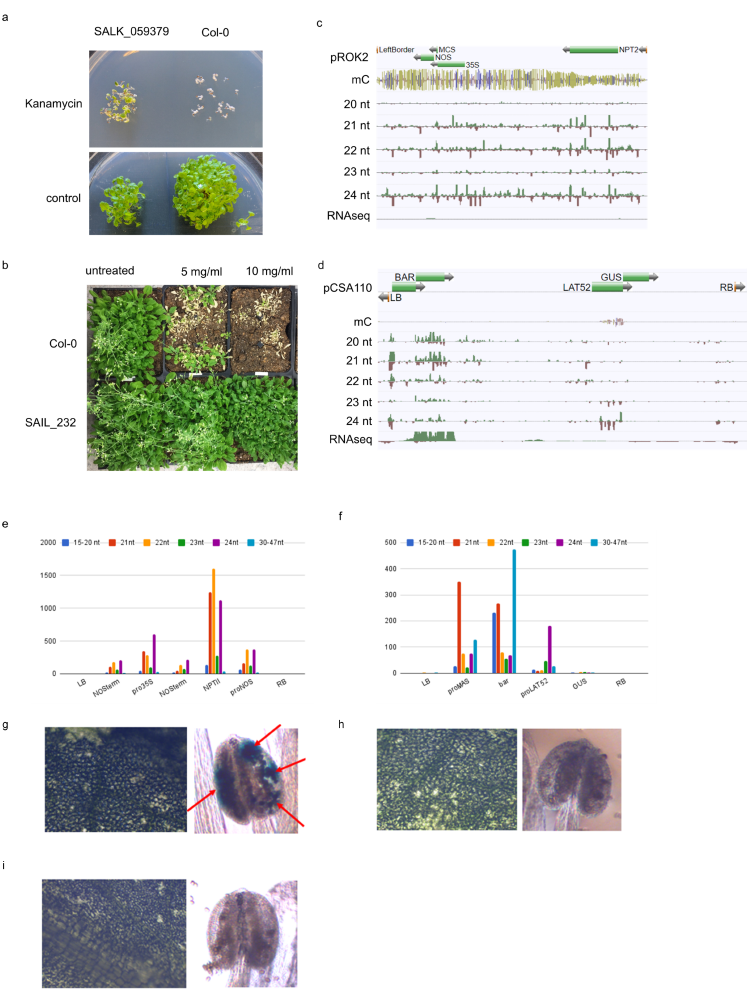
SALK and SAIL T-strand sequences show divergent epigenomic signatures. Transgene activity was tested by exposing plants to the respective selective agent: (a) SALK_059379 grown on media containing the antibiotic kanamycin (50ug/uL) (or empty control) and (b) the SAIL_232 sprayed with the herbicide Finale™ at two different concentrations through the middle of the tray. All phenotypes are compared to the WT Col-0 (a,b). Analysis of expression and epigenetic signatures on the corresponding T-DNA sequence is captured in browser shots for SALK_059379 plasmid pROK2 (c) and SAIL_232 plasmid pCSA110 (d): Illumina read mapping of bisulfite sequencing, RNA-seq and different small RNA (sRNA) species. Quantification of individual siRNA read length against individual parts of the two plasmid sequences are reported in (e = pROK2) and (f = pCSA110). GUS staining of leaves and flowers of SAIL_232 (g), with Col-0 (h) and SALK_059379 (i) as control.

### Differential siRNA species define transgene silencing

We hypothesized that the observed plasmid dependent transgene expression is dictated by gene silencing via RdDM pathways [35]. To test for the presence of small RNA (sRNA) and cytosine DNA methylation within T-DNA loci, we sequenced sRNA populations, as well as bisulfite-converted whole genome DNA libraries for both lines. Our analyses identified abundant, yet divergent sRNA species (15-20nt, 21nt, 22nt, 23nt, 24nt) that mapped to genomic insertions of the *nptII* (pROK2) and *bar* (pCSA110) vector sequences in the SALK and SAIL lines, respectively (Fig. 3c,d). The *nptII* gene was highly targeted by 21/22nt short interfering (si)RNAs, as well as RdDM promoting 24nt siRNAs. The NOSp and duplicated NOSt were equally targeted by 22nt and 24nt siRNAs, although at lower numbers than the *nptII* gene itself (Fig. 3e). In contrast, the SAIL_232 *bar* gene and its promoter were targeted by high numbers of 21nt siRNAs, and lacking 24-mers (Fig. 3f). Interestingly, the pCSA110 encoded pollen specific promoter pLAT52, promoting the reporter gene *GUS*, was the only element highly targeted by 24nt siRNAs as well as high cytosine methylation levels in all sequence contexts (CG, CHG and CHH methylation, where H is A, C or T) in the SAIL background (Fig. 3d, f; Supplementary Table 6). In contrast, the entire pROK2 T-DNA region showed high cytosine methylation in all three contexts. We found all elements to be fully CG and CHG methylated, and between 30% (*nptII*) and 71% (NOSt) of CHH’s were methylated (Supplementary Table 7). In summary, the *GUS* reporter in pCSA110 and all pROK2 transgenic elements are targeted by RdDM gene silencing pathway 24nt siRNAs. In contrast, the SAIL_232 pCSA110 *bar* gene was only targeted by 21/22nt siRNAs without any apparent effects on expression or the chemical resistance phenotype.

We were curious whether the 24nt siRNA and DNA methylation of the *GUS* gene observed in leaf tissue suppresses expression in pollen, where it is driven by the strong pLAT52 promoter. Indeed staining of mature pollen identified GUS activity in the tested SAIL line, thus suggesting that these epigenetic effects can be partially overcome by when transgene expression is driven by a strong promoter (Fig. 3g-i).

While these observations are limited to the transformation vectors, we specifically looked at the individual junctions between genome and T-DNA to examine for epigenetic effects on the flanking genomic DNA sequences/genes. We only found few siRNA reads at two SALK_059379 (of eight) and six SAIL_232 (of 11), and thus were not able to draw any conclusions. Similarly for bisulfite-converted DNA reads, where we were able to identify DNA methylation signatures at only one junction in each transgenic line. With the caveat that only a few cases were examined, we were unable to observe any signatures for siRNA or DNA methylation spreading outside of the T-DNA borders.

### T-DNA-insertions shape the local chromatin environment

Besides DNA methylation, modifications of histone tails as well as histone variants are also key epigenome features playing crucial roles in instructing gene expression dynamics [36]. To study the role of T-DNA insertions in shaping the adjacent chromatin in SALK_059379 and SAIL_232 seedlings, we employed ChIP-seq assays to profile the occupancy of two histone modifications (H3K4me3, H3K27me3) and the histone variant H2A.Z (http://neomorph.salk.edu/T-DNA_ChIP.php). H3K4me3 marks active or poised genes [37], whereas H3K27me3 is involved in Polycomb Repressive Complex 2 (PRC2)-mediated gene silencing [38]. The histone variant H2A.Z is known to be involved in the gene responsiveness to environmental stimuli in *Arabidopsis* [39]. SALK_059379 carries five T-DNA-insertions, two of which are homozygous and could thus be analyzed and visualized in our ChIP-seq data (Fig. 4a and Supplementary Fig. 5a-c). The SALK:chr2_18Mb T-DNA insertion led to a 5.5 kb deletion ranging from the promoter of *At2g45820* to the gene body of *At2g45840* (Fig. 1d, 4a). We observed that the H3K4me3 domain at *At2g45820* (domain 1) is affected by the deletion, displaying lower levels of H3K4me3 (Fig. 4a and Supplementary Fig. 5a). In Col-0, this deleted fragment shows a large H3K27me3 domain which extends into the gene body of *At2g45840*. Levels of H3K27me3 are also reduced at that region (domain 2) in SALK_059379 (Fig. 4a and Supplementary Fig. 5a). The SALK:chr2_15Mb insertion occurred in the first intron of *At2g36290*, and leads to a strong reduction of H3K4me3 and H2A.Z compared to Col-0 (Supplementary Fig. 5b, c).

**Fig. 4:**
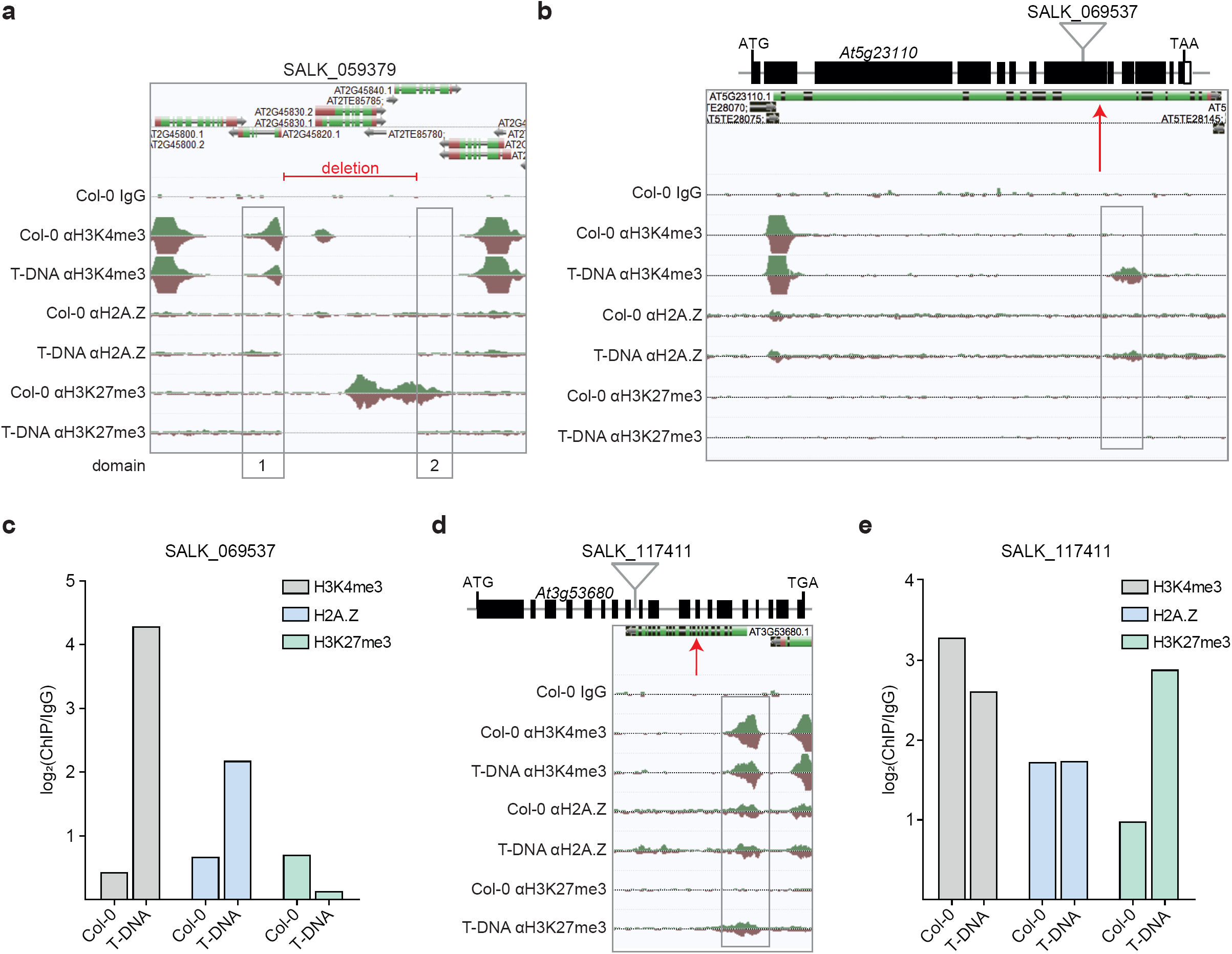
T-DNA insertions alter the chromatin landscape. a) AnnoJ genome browser visualization of ChIP-seq reads derived from H3K4me3, H2A.Z and H3K27me3 occupancy in Col-0 and SALK_059379 seedlings around the T-DNA-induced deletion on the chromosome 2. Histone domains that are adjacent to the T-DNA deletion are indicated as domain 1 and 2. (b) Annoj genome browser visualization of H3K4me3, H2A.Z and H3K27me3 occupancy in Col-0 and SALK_069537 seedlings around the T-DNA insertion at *At5g23110*. Schematic illustration shows the gene structure of *At5g23110* and the approximate location of the corresponding T-DNA insertion. (c) Quantification of H3K4me3, H2A.Z and H3K27me3 around the T-DNA insertion site at *At5g23110* (SALK_069537). (d) Visualization of H3K4me3, H2A.Z and H3K27me3 occupancy in Col-0 and SALK_117411 seedlings indicates *de novo* trimethylation of H3K27 around the T-DNA insertion in *At3g53680*. Scheme of gene structure of *At3g53680* shows the approximate location of the T-DNA insertion in genome of SALK_117411 plants. (b) Levels of H3K4me3, H2A.Z and H3K27me3 around the T-DNA insertion at *At3g53680* in SALK_117411 seedlings are shown. A red arrow marks the approximate location of the T-DNA integration site in the Annoj genome browser screenshots. The respective occupancies were identified with ChIP-seq and all shown AnnoJ genome browser tracks were normalized to the respective sequencing depth. The Col-0 IgG track serves as a control. The occupancy of the respective domains was calculated as the ratio between the respective ChIP-seq sample and the Col-0 IgG control.

For SAIL_232, we began analysis of the chromatin environment at the Col-3 specific large inversion on chromosome 1. The most profound impact at the flanking sites was found on the level of H3K27me3, where a large H3K27me3 domain, that spans *At1g32420* and *At1g32430* in Col-0 seedlings, was split and yet still maintained at the new inverted chromosomal positions in Col-3 (Supplementary Fig. 5d, e). As a result, we discovered a new H3K27me3 domain (domain 2) adjacent to *At1g33700* that is likely caused by spreading of H3K27me3 from *At1g32420* into the inverted genomic region (Supplementary Fig. 5d, e). In contrast, the H3K27me3 domain at the gene body of *At1g32430* was detached from the larger H3K27me3 domain and decreased in size compared to Col-0, possibly due to the loss of H3K27me3 reinforcement (Supplementary Fig. 5d, e). The T-DNA insertion in SAIL:chr5_15Mb occurred in the first exon of *At5g38850*, and changed the H3K4me3 pattern in this genomic area (Supplementary Fig. 5f, g). Levels of H3K4me3 were clearly lower at *At5g38850* and started to spread into the adjacent *At5g38840* gene (Supplementary Fig. 5f, g).

To further our understanding of how the insertion of T-DNA affects the adjacent chromatin environment, we expanded our analysis to include eight additional T-DNA integration events (http://neomorph.salk.edu/T-DNA_ChIP.php), previously discovered in five T-DNA lines (SALK_061267, SALK_017723, SALK_069537, 50B_HR80 and SALK_117411) (Supplementary Table 8). Initially, we focused on two single exon genes: *At1g32640* (SALK_061267) and *At5g11930* (SALK_017723). In Col-0 WT *At1g32640* carries H3K4me3 and H2A.Z over the entire gene body whereas levels of H3K27me3 were comparably low (Supplementary Fig. 5h, i). The T-DNA insertion at *At1g32640* disrupts the H3K4me3 domain, leading to an overall reduction, and also disrupts the H2A.Z domain resulting in a clear increase, where especially the first domain shows a 3-fold increase (Supplementary Fig. 5h, i). At an additional T-DNA insertion site, 865 bp outside of the gene body of *At1g23910*, we observed a decrease in all three gene-body localized marks (Supplementary Fig. 5j, k). We discovered an even stronger impact at *At5g11930* in SALK_017723 where the gene body T-DNA insertion caused a complete loss all three profiled histone modifications/variants (Supplementary Fig. 6a, b).

Next, we analyzed T-DNA insertions in multi-exon genes. Contrary to *At1g32640* and *At5g11930* where H3K4me3 and H2A.Z can be detected over the entire gene body in Col-0 seedlings (Supplementary Fig. 5h, 6a), chromatin marks are restricted to the +1 and +2 nucleosome region at *At5g23110* (SALK_069537) (Fig. 4b). Strikingly, SALK_069537 seedlings display a new H3K4me3 and H2A.Z domain downstream of the T-DNA insertion in *At5g23110* which is not present in Col-0 seedlings (Fig. 4b, c). This is the first evidence of a T-DNA-induced *de novo* tri-methylation of H3K4 and incorporation of H2A.Z in the host genome.

In order to examine whether this pattern was specific to *At5g23110*, we inspected another T-DNA line with a T-DNA insertion in *At3g57300*, another large multi-exon gene (Supplementary Fig. 6c, d). This 50B_HR80 T-DNA line is derived from a T-DNA mutant-based screen for defects in homologous recombination [40]. Similar to SALK_061267, 50B_HR80 also shows the *de novo* H3K4 trimethylation and H2A.Z incorporation adjacent to the T-DNA insertion site (Supplementary Fig. 6c, d), suggesting that a more general mechanism may takes place at T-DNA insertions within larger genes.

In addition, we also found evidence of *de novo* trimethylation of H3K27 next to a T-DNA insertion site in the gene body of *At3g53680* in SALK_117411 seedlings (Fig. 4d, e). This new H3K27me3 domain expands into the promoter region of *At3g53680* and affects H3K4me3 and H2A.Z only slightly (Fig. 4d, e). SALK_117411 carries another T-DNA in the 3’UTR of *At1g01700* which has a very selective impact on the chromatin environment at *At1g01700* reducing only H3K27me3 whereas H3K4me3 and H2A.Z remained unaffected (Supplementary Fig. 6e, f). Another 3’UTR-located T-DNA insertion in *At3g46650* (Supplementary Fig. 6g) also impacted the adjacent chromatin in SALK_069537 but this time H3K4me3 and H2A.Z and not H3K27me3 were profoundly affected (Supplementary Fig. 6g, h). Taken together, our ChIP-seq assays clearly demonstrate that T-DNA integration affects the local chromatin environment at the site of insertion.

## Discussion

During the process of Agrobacterium transformation, T-DNA can insert itself into the plant genome at random or induced (e.g. through nuclease activity) sites of chromosomal double-strand breaks, utilizing host DNA repair mechanisms. The ideal transgene delivery system would result in the gene of interest being integrated as a single functional copy. Unfortunately, the majority of T-DNA transformed plants deviate from this ideal (reviewed by Gelvin [5]), containing only partially integrated or concatemers of T-DNA that often are not expressed (silenced) (e.g. Gelvin [10], Peach and Velten [41]). The genome structure at sites of T-DNA insertion or the structure of the T-DNA insertion itself can additionally induce epigenetic alterations with detrimental effects on transgene function. To better understand these effects, short-read sequencing technologies have been employed [42]. Unfortunately, the repetitiveness of concatenated T-DNAs (carrying transgene sequences) and vector backbone insertions [13] has hampered efforts to fully appreciate the complexity of these events and their impact on the plant genome and epigenome.

Using a combination of long-read sequencing/mapping tools, we identified and analyzed perturbations to the genome structure of four randomly selected transgenic, floral dip transformed *Arabidopsis thaliana* (Columbia accession) T-DNA insertion lines from three of the most widely utilized plant mutant collections (SALK, SAIL, WISC). Optical genome physical maps (BNG) created from single DNA molecules with an average size of up to 288 kb were critical to unveil both the size and structure of genomic transgene insertions which ranged in size up to 236 kb in length. Using nanopore long read DNA sequencing technology we were subsequently able to assemble contigs up to chromosome arm length, which closed 83% of non-centromeric (26% of all) misassemblies within the gold-standard reference genome (TAIR10) [43]. Sequence contigs for the two transgenic lines (SAIL_232 and SALK_059379) captured up to 39 kb of assembled T-DNA insertion sequence and revealed the complexity of *Agrobacterium*-mediated transgene insertions. Less repetitive rearrangements, like the SAIL-specific 512 kb inversion on chromosome 1, were perfectly captured within single chromosome arm spanning contigs. Although ONT long read sequences were not sufficient to provide complete contiguity of the highly repetitive T-DNA insertions, BNG maps preserved contiguity over all insertions and rearrangements, and thus provided absolute proof for these events - in contrast to e.g. split-read mapping of short Illumina reads [42].

We hypothesize that the final T-DNA insertion derives from two independent T-DNA insertions at each end of the genomic breakpoint, subsequent connection into the observed long T-DNA concatemers, and possible interaction among the many resulting homologous regions. Previously these long repetitive segments were attributed to the floral dip transformation method Arabidopsis [22]. Long homologous stretches between the independently inserted T-DNA/vector backbone concatemers represent inverted repeats which can lead to hairpin formation, which stalls the recombination fork and excision during DNA repair [45]. We identified a single chimeric vector/vector insertion (SALK:chr2_15Mb) that most likely occurred through homology-dependent recombination of the two identical NOSt sequences in the T-DNA vector (pROK2:6778-7038 bp and 8607-8867 bp). While internal breakpoints and the scrambled insertion pattern that we observed are in support of this, we have no data to confirm whether this happened before, during, or after integration (Fig. 1, 2, Supplementary Fig. 3). Further experiments are necessary to characterize the mechanism of T-DNA strand concatenation, the role of floral dip and whether internal rearrangements and excisions occur during the insertion process or at later meiotic stages.

The relationship between *Agrobacterium*-mediated T-DNA integration and the host epigenome are largely uncharted territory. Reports and anecdotal experiences suggested silencing of the kanamycin antibiotic resistance *nptII* marker gene in a large number of SALK lines [4]. In contrast silencing of the SAIL line inherent herbicide tolerance *bar* gene has not been reported. Our analyses revealed distinct epigenomic features for the two phenotypically indistinct transgenic lines; both transgenes are targeted for sRNA-level silencing. For pROK2, the sRNAs have progressed to the RdDM phase, and are targeting the entire transgene for methylation and functional shut-down of the resistance gene. It appears that the antibiotic resistance marker *nptII* is undergoing a mixture of expression-dependent silencing and conventional Pol IV-RdDM. It lacks heavy 24mers at the promoter and makes 21-22mers within the coding region. These even levels of 21-22 and 24nt siRNAs further suggest that the *nptII* gene is transcribed at least at some level but degraded very efficiently and targeted for RdDM. This transgene is a heavy target of some type of RdDM (equal levels of CHH and CG) and produces no steady state mRNA, confirming the antibiotic sensitive phenotype.

For pCSA110, besides the Lat52 promoter, the *bar* transcript is targeted by RNAi, likely RDR6-dependent since 21mers are derived from both strands. The few observed 24nt siRNA traces are potentially degradation products from the highly expressed *bar* gene. However, the siRNAs are not successfully targeting RdDM, most likely due to absence of Pol V expression, so there is likely a reduced level of steady state mRNA, but enough to make a protein and provide the observed herbicide tolerance. It is not surprising that the pollen LAT52 promoter is heavily targeted by Pol IV-RdDM. This tomato-derived promoter is made up of retrotransposon sequences that show sufficient sequence conservation among plant species, that this promoter is always recognized and targeted by Pol IV-RdDM. We were able to show that this strong promoter can override the silencing machinery to drive *GUS* expression in mature pollen.

Our ChIP-seq-based analyses of H3K4me3, H3K27me3 and H2A.Z occupancy surrounding ten T-DNA insertion sites and one deletion site revealed a strong impact of T-DNA insertion on the local chromatin environment. The most profound effects were found at T-DNA insertions in large multi-exonic genes which gave rise to completely new H3K4me3/H2A.Z domains. From other systems such as yeast or humans it is known that the histone variant H2A.Z as well as the histone modification H3K4me3 are involved in the DNA damage response, both rapidly accumulating at double strand breaks [51, 52]. Interestingly the T-DNA integration in SALK_069537 and 50B_80HR T-DNA lines results in only one new domain at the 3’ side of the gene, clearly arguing against a DNA damage response scenario, suggesting that the direction of transcription spatially determines the T-DNA insertion-facilitated *de novo* histone domain formation. Moreover, we also discovered a T-DNA insertion-induced *de novo* trimethylation of H3K27 spanning from the site of insertion into the promoter. The asymmetric shape of the domain with its gradual increase towards the promoter suggests that it is caused by the *de novo* recruitment of PRC2 and/or the blocking of H3K27me3 demethylases such as REF6 [50] in that region and not by PRC2-mediated spreading of H3K27me3 from the T-DNA. Our chromatin study identified a highly diverse and complex impact of T-DNA insertions on the adjacent chromatin environment. Future studies will need to be conducted to analyze how T-DNA insertions shape the local host epigenome.

Its small genome size and the widespread utilization of the T-DNA mutant collections made *Arabidopsis thaliana* an ideal organism to study the structural and transgenerational effects of transgene insertions. Our findings pave the way for structural genomic studies of transgenic crop plants and provide insights into the effects of transgene insertions on the epigenetic landscape.

## Material and Methods

### Plant Material

Seeds were ordered from ABRC (https://abrc.osu.edu): SALK_059379 (segregant and homozygous lines), SALK_075892, SAIL_232 (seg.), SAIL_59 (seg.), SAIL_107 (seg.), WiscDsLox_449D11 (seg.), SALK_061267, SALK_017723, SALK_069537 and SALK_117411. 50B_HR80 seeds were kindly provided from Susan M. Gasser. Plants were grown in a 20⁏ growth room with 13h light/11h dark cycles in peat-based soil supplemented with nutrients, and collected approximately three weeks after bolting. Growth conditions for plant material that was used for ChIP-seq is listed in Supplementary Table 8.

### Testing Selection Markers

#### Kanamycin resistant lines- SALK and Wisc

MS/MES media was prepared (1L NP H2O, 2.2g MS, 0.5g MES, pH5.7) and autoclave sterilized. Kanamycin (50ug/mL) was added to half of the media, and plates with and without the antibiotic were poured. Seeds from the lines SALK_059379 (homozygous), SAIL_232, WiscDsLox_449D11, and Col-0 CS70000 control were sterilized and spotted on the plates, which were placed in the same growth conditions as the plant material in soil. Survival rates were measured 5 days after seeds were spotted and growth was observed for further 14 days.

#### Finale™ *resistant lines- SAIL*

One flat each of Col-0 (control) and SAIL_232 were grown in soil as outlined in *Plant Material*. Plants were sprayed with Finale™ at three different concentrations (5mg/L, 10mg/L, 20mg/L) at 5, 14, and 21 days after germination [1, 2].

#### GUS reporter staining- SALK and SAIL

Leaf and flower tissue was submerged in 1.5mL staining buffer (0.5mM ferrocyanide, 0.5mM ferricyanide, 0.5% Triton, 1mg/mL X-Gluc, 100mM Sodium Phosphate Buffer pH 7) and incubated at 37°C overnight. A light microscope was used to visualize the staining at 10x magnification for the leaves and 4x for the flowers.

### ONT Library Prep and Sequencing

5g of flash-frozen leaf tissue, pooled from the segregant seed stocks, was ground in liquid nitrogen and extracted with 20mL CTAB/Carlson lysis buffer (100mM Tris-HCl, 2% CTAB, 1.4M NaCl, 20mM EDTA, pH 8.0) containing 20μg/mL proteinase K for 20 minutes at 55⁏. To purify the DNA, 0.5x volume chloroform was added, mixed by inversion, and centrifuged for 30 minutes at 3000 RCF. Purification was followed by a 1x volume 1:1 phenol: [24:1 chloroform:isoamyl alcohol] extraction. The DNA was further purified by ethanol precipitation (1/10 volume 3M sodium acetate pH 5.3, 2.5 volumes 100% ethanol) for 30 minutes on ice. The resulting pellet was washed with freshly-prepared ice-cold 70% ethanol, dried, and resuspended in 350μL 1x TE buffer (10mM Tris-HCl, 1mM EDTA, pH 8.0) with 5μL RNase A (Qiagen, Hilden) at 37⁏ for 30 minutes, then at 4⁏ overnight. The RNase A was removed by double extraction with 24:1 chloroform:isoamyl alcohol, centrifuging at 22,600x*g* for 20 minutes at 4⁏ each time to pellet. An ethanol precipitation was performed as before for 3 hours at 4⁏, washed, and resuspended overnight in 350μL 1x TE buffer. The genomic DNA samples were purified with the Zymogen Genomic DNA Clean and Concentrator-10 column (Zymo Research, Irvine). The purified DNA was prepared for sequencing with the Ligation Sequencing Kit 1D (SQK-LSK108, ONT, Oxford, UK) sequencing kit protocol. Briefly, approximately 2 μg of purified DNA was repaired with NEBNext FFPE Repair Mix for 60 min at 20°C. The DNA was purified with 0.5X Ampure XP beads (Beckman Coulter, Brea). The repaired DNA was End Prepped with NEBNExt Ultra II End-repair/dA tail module and purified with 0.5X Ampure XP beads. Adapter mix (ONT, Oxford, UK) was added to the purified DNA along with Blunt/TA Ligase Master Mix (NEB) and incubated at 20°C for 30 min followed by 10 min at 65°C. Ampure XP beads and ABB wash buffer (ONT, Oxford, UK) were used to purify the library molecules, which were recovered in Elution buffer (ONT, Oxford, UK). The purified library was combined with RBF (ONT, Oxford, UK) and Library Loading Beads (ONT, Oxford, UK) and loaded onto a primed R9.4 Spot-On Flow cell. Sequencing was performed with an ONT MinION Mk1B sequencer running for 48 hrs. Resulting FAST5 (HDF5) files were base-called using the ONT Albacore software (0.8.4) for the SQK-LSK108 library type.

### Sequence extraction, assembly, consensus and correction

Raw ONT reads (fastq) were extracted from base-called FAST5 files using poretools [3]. Overlaps were generated using minimap [4] with the recommended parameters (-Sw5 -L100 - m0). Genome assembly graphs (GFA) were generated using miniasm [4]. Unitig sequences were extracted from GFA files. Three rounds of consensus correction was performed using Racon [5] based on minimap overlaps, and the resulting assembly was polished using Illumina PCR-free 2×250 bp reads mapped with bwa [6] and pilon [7]. Genome statistics were generated using QUAST [8].

The ONT reads were also assembled with the CANU assembler [9]. However, the minimap/miniasm assemblies were of higher contiguity and resolved longer stretches of the T-DNA insertions. In addition, the minimap/miniasm assemblies were computed within 4-6 hours, while the CANU assemblies took between one (Col-0) and four weeks (SAIL). We hypothesize that the SAIL lines inherently complex repeat structure caused the long compute time, while the reference Col-0 required only the expected time of one week. Also, we hypothesize that minimap/miniasm resolved the T-DNA structures more fully due to the fact it does not have a read correction step, which could lead to the collapsing of highly repetitive yet distinct T-DNA insertions.

### Identification of individual T-DNA matching reads and annotation

Reads obtained via ONT sequencing were imported into Geneious R10 [10], along with the pROK2, and pCSA110 plasmid sequences. BLAST databases, were created for the complete set of SALK_059379 and SAIL_232 ONT reads, and their respective plasmid sequences were BLASTn searched against these databases. Reads with hits on the plasmid sequences were extracted, and BLASTn searched against the TAIR10 reference. NCBI megaBLAST [11] and Geneious ‘map to reference’ functions were used to annotate regions on each read corresponding to either TAIR10 or the plasmid sequence. The chromosomal regions were compared to the regions identified by T-DNA seq and BNG mapping to verify and refine T-DNA insertion start and stop coordinates.

### BNG Optical Genome Mapping

High molecular weight (HMW) DNA for BNG mapping and ONT sequencing was extracted as outlined in Kawakatsu [12]. Briefly, up to 5g of fresh mixed leaf and flower tissue (excluding chlorotic leaves and stems) pooled from the segregant seed stocks were homogenized in 50mL nuclei isolation buffer. Nuclei were separated from debris using a Percoll layer. Extracted nuclei were subsequently embedded in low melting agarose plugs and exposed to lysis buffer overnight. DNA was released by digesting the agarose with Agarase enzyme (Thermo Fisher Scientific).

HMW DNA was nicked with the enzyme Nt.BspQI (New England Biolabs, Ipswich, MA), fluorescently labeled, repaired and stained overnight following the Bionano Genomics nick-labeling protocol and accompanying reagents (Bionano Genomics, San Diego, CA) [13]. Each *Arabidopsis* T-DNA insertion line was run for up to 120 cycles on a single flow cell on the Irys platform (Bionano Genomics, San Diego, CA). Collected data was filtered (SNR = 2.75; min length 100kb) using IrysView 2.5.1 software, and assembled using default “small” parameters. Average molecule length for assembly varied between 201 kb and 288 kb, and resulted in BNG map N50 0.97 to 1.03 Mb. Derived assembled maps were anchored to converted TAIR10 chromosomes using the RefAligner tool and standardized parameters (-maxthreads 32 -output-veto-filter _intervals.txt$ -res 2.9 -FP 0.6 -FN 0.06 -sf 0.20 -sd 0.0 -sr 0.01 -extend 1 –outlier 0.0001 -endoutlier 0.01 -PVendoutlier -deltaX 12 -deltaY 12 -xmapchim 12 -hashgen 5 7 2.4 1.5 0.05 5.0 1 1 1 -hash -hashdelta 50 -mres 1e-3 -hashMultiMatch 100 -insertThreads 4 -nosplit 2 - biaswt 0 -T 1e-10 -S -1000 -indel -PVres 2 -rres 0.9 -MaxSE 0.5 -HSDrange 1.0 -outlierBC - xmapUnique 12 -AlignRes 2 -outlierExtend 12 24 -Kmax 12 -f -maxmem 128 -stdout -stderr). We utilized the resulting *.xmap, *_q.cmap and *_r.cmap files in the structomeIndel.py script (https://github.com/RyanONeil/structome) to identify T-DNA insertion locations and sizes. We used this script to further determine misassemblies in the ONT genome assemblies. Known Col-0 mis-assemblies [12] were subtracted from the list of derived locations.

### Aligning contigs to TAIR10

We aligned ONT contigs to the TAIR10 reference using the BioNano Genomics RefAligner as above (Supplementary Table 3). ONT contigs were in silico digested with Nt.BspQI and used as template in RefAligner. The alignment output was manually derived from the.xmap output file.

### Genotyping of structural genome variations

Primer3 was used to create oligos (Supplementary Table 5) for each insertion identified in the SALK_059379 and SAIL_232 lines. The forward primers correspond to the chromosome sequence ∼500bp before the insertion start site, and the reverse to the plasmid sequence ∼500bp after the insertion start site. A second set of reverse primers corresponding to the original WT chromosome ∼1kb after the original forward primers. Individuals and pooled tissue from SALK_059379, SAIL_232, SAIL_59, SAIL_107 and Col_0 CS7000 DNA was extracted using Qiagen DNeasy Plant Mini kit (following the protocol) and genotyped using these primers, with Col-0 as the negative control.

### Bisulfite Sequencing

DNA was extracted using the DNeasy Plant Mini Kit (Qiagen, Hilden, D) according to the manufacturer’s protocol from segregant plant pools, and quantified using the Qubit dsDNA BR assay kit. Illumina sequencing library preparation and bisulfite conversion was conducted as described in Kawakatsu [12]. Briefly, DNA was End Repaired using the End-It kit (Epicentre, Madison, WI), A-tailed using dA-Tailing buffer and 3μL Klenow (3’ to 5’ exo minus) (NEB, Ipswich, MA) and Truseq Indexed Adapters (Illumina, San Diego, CA) were ligated overnight. Bead purified DNA was quantified using the Qubit dsDNA BR assay kit, and stored at −20⁏. At least 450ng of adapter ligated DNA was taken into bisulfite conversion which was performed according to the protocol provided with the MethylCode Bisulfite Conversion kit (Thermo Fisher, Waltham, MA). Cleaned, converted single stranded DNA was amplified by PCR using the KAPA U+ 2x Readymix (Roche Holding AG, Basel, CH): 2 min at 95⁏, 30 sec at 98⁏ [15 sec at 98⁏, 30 sec at 60⁏, and 4 min at 72⁏] x 9, 10 min at 72⁏, hold at 4⁏. After amplification, the DNA was bead purified, the concentration was assessed using the Qubit dsDNA BR assay kit, and the samples were stored at −20⁏. WGBS library was sequenced as part of a large multiplexed pool paired-end 150 bp on an Illumina HiSeq 4000.

### RNA Libraries

RNA extractions performed using the RNeasy Plant Mini Kit (Qiagen, Hilden, D) from segregant plant pools. RNA-seq libraries were prepared manually (SAIL_232) as described in Kawakatsu or using the Illumina NeoPrep Library Prep System (SALK_059379) (Illumina, San Diego, CA), following the NeoPrep control software protocol. The completed libraries were quantified using the Qubit dsDNA HS assay kit and stored at −20⁏. SALK_059379 was sequenced in duplicate as single end 150 bp in a multiplexed pool on two lanes of an Illumina HiSeq 2500. SAIL_232 was sequenced as part of a large multiplexed pool paired-end 150 bp on an Illumina HiSeq 4000.

### small RNA Libraries

We have extracted sRNA according to the protocol described in Vandivier [14] with modifications to the RNA extraction and size selection steps. In brief, 300mg flash-frozen leaf tissue from segregant plant pools was ground in liquid nitrogen and extracted with 700μL QIAzol lysis reagent (Qiagen, Hilden, D). The RNA was separated from the lysate using QIAshredder columns (Qiagen, Hilden, D) and purified with a ⁏ volume 24:1 chloroform:isoamyl alcohol (Sigma-Aldrich, St. Louis, MO) extraction followed by a 1mL wash of the aqueous phase with 100% ethanol. The RNA was further purified using miRNeasy columns (Qiagen, Hilden, D) with two 82μL DEPC-treated water washes. 37μL DNase was added to the flow-through and incubated at RT for 25 minutes. The DNase-treated RNA was precipitated with 20μL 3M sodium acetate (pH 5.5) and 750μL 100% ethanol overnight at −80⁏. The pellet was washed with 750μL ice-cold 80% ethanol and resuspended in a 12:1 DEPC-treated water:RNase OUT Recombinant Ribonuclease Inhibitor (Thermo Fisher, Waltham, MA) mixture on ice for 30 minutes and quantified (Qubit RNA HS assay kit).

Size selection and library prep were conducted exactly as described in Vandivier [14]. The RNA from 15-35bp was cut out of the gel and purified. sRNA libraries were sequenced in duplicate as single end 150 bp in a multiplexed pool on two lanes of an Illumina HiSeq 2500.

### Short read analysis

RNA-seq and sRNA Illumina reads were adapter trimmed and mapped against the junction sequences as well as the corresponding plasmid sequences, using Bowtie2 and Geneious aligner (Geneious R10.2.3; custom sensitivity, iterate up to 5 times, zero mismatches or gaps per read, word length 14) at highest stringency. RNA-seq reads were mapped against AtACT1 as control, and found this gene expressed. RNA-seq reads were mapped against Pol V AT2G40030 as control, and found this gene not expressed. Resulting reads were extracted and re-mapped against the TAIR10 reference genome to ensure that no off-target reads mapped to the plasmid sequences. Because we analyzed various genomic insertions against a single reference, we did not normalize read mapping.

The MethylC-Seq data (paired-end) reads in FastQ format and the second pair reads were converted to their reverse complementary strand nucleotides. Then reads were aligned to the *Arabidopsis thaliana* reference genome (araport 11) and pCSA110/pROK2 genome. The Chloroplast genome was used for quality control (< 0.35% non-conversion rate). After mapping, overlapping bases in the paired-end reads were trimmed. The base calls per reference position on each strand were used to identify methylated cytosines at 1% FDR.

### ChIP-seq assays and analysis

ChIP-seq experiments were performed as previously described [15] using antibodies against H3K4me3 (04-745, EMD Millipore), H2A.Z (39647, Active Motif) and H3K27me3 (39156, Active Motif). Incubation with mouse IgG (015-000-003, Jackson ImmunoResearch) served as our negative control. ChIP DNA was used to generate sequencing libraries following the manufacturer’s instructions (Illumina). Libraries were sequenced on the Illumina Genome Analyzer II. Sequencing reads were aligned to the TAIR10 genome assembly using Bowtie2 [16]. Histone domains next to the T-DNA insertion events were identified with SICER [17]. The bigWigAverageOverBed tool executable from the UCSC genome browser [18] was used to quantify the occupancy of H3K4me3, H2A.Z and H3K27me3 next to T-DNA integration sites. The TAIR10 reference domain coordinates can be found in Supplementary Table 9.

## Data Availability

Raw ONT sequencing data was deposited in the European Nucleotide Archive (ENA) under project PRJEB23977 (ERP105765). Final polished assemblies were deposited in the ENA Genome Assembly Database PRJEB23977. Raw BNG molecules and assembled maps are deposited under BioProjects PRJNA387199, PRJNA387199, PRJNA387199 and PRJNA387199. Short-read datasets are deposited under GEO accession GSE108401.

## Author contributions

FJ, TPM, JRE conceptualized the study; AR, FJ, STM, MZ, RC, JN and JS created sequencing libraries and/or conducted sequencing; FJ, AR, HC, MZ and TPM analyzed sequencing data; AR, JS and FJ conducted plant studies; MZ conceptualized and conducted ChIP-seq experiments; FJ, AR, TPM, RKS, MZ and JRE wrote the manuscript.

## Acknowledgments

We would like to thank Cesar Barragan and Dr. Ronan O’Malley for insights into the T-DNA project, and Bruce Jow and Christopher Santos for excellent greenhouse support. We thank Detlef Weigel and Christa Lanz, both Max Planck Institute for Developmental Biology (Tuebingen, Germany) for performing and providing Illumina short read sequencing for Col-0 CS70000. FJ was supported through a Human Frontier Science Program Organization long-term fellowship. M.Z. was supported by the Salk Pioneer Postdoctoral Endowment Fund as well as by a Deutsche Forschungsgemeinschaft (DFG) research fellowship (Za-730/1-1). JRE is an Investigator of the Howard Hughes Medical Institute.

---

**Supplementary Table 1: Bionano Genomics and Oxford Nanopore Technologies sequencing and assembly statistics.**

**Supplementary Table 2: Genome Alignment of ONT assembled contigs against TAIR10 using the Bionano Genomics RefAligner algorithm.**

**Supplementary Table 3: Identification of ‘N’ regions in the TAIR10 reference genome and the corresponding Col-0 ONT contigs.**

**Supplementary Table 4: BNG maps identify misassembled regions in ONT contigs as ‘False Duplications’ or ‘False Deletions’.**

**Supplementary Table 5: List and analysis of genomic T-DNA insertion sites.**

**Supplementary Table 6: Analysis of methylated cytosines on the pCSA110 plasmid for SAIL_232.**

**Supplementary Table 7: Analysis of methylated cytosines on the pROK2 plasmid for SALK_059379.**

**Supplementary Table 8: Experimental information for conducted ChIP experiments in T-DNA and WT Arabidopsis lines.**

**Supplementary Table 9: TAIR10 reference histone domain coordinates.**

**Supplementary Table 10: PCR oligo sequences used for genotyping T-DNA inserts**

**Supplementary Fig. 1:** Alignment of *de novo* assembled ONT contigs for Col-0 (CS70000) and the transgenic lines SALK_059379 and SAIL_232 relative to the reference TAIR10. Uninterrupted colored blocks indicate contig length, black bars indicate the start and end of contigs, black boxes indicate centromere gaps. Arrowheads represent T-DNA insertion sites; orange lines and boxes indicate sites of translocations. Drawn to scale for each chromosome individually.

**Supplementary Fig. 2: Background inversion in SAIL lines derived from Col-3.** Two additional SAIL lines were chosen at random and genotyped along with SAIL_232. The primers were designed to amplify 1000bp spanning the junction between the genomic and inverted DNA sequence (based on *de novo* reads from ONT sequencing). The inversion on chromosome 1 was seen in all SAIL lines tested, unlike the other chromosomal architecture disruptions found. Thus, it can be concluded that this inversion is part of the Col-3 background from which SAIL lines are derived. The bands at 1500bp in the SAIL_107 and SAIL_59 lines are off-target amplification products.

**Supplementary Fig. 3:** Model for microhomology dependent excision of T-DNA/genome fragments. (a) A double strand chromosome break leads to (b) two multi-copy T-DNA strand insertions in opposing directions. The exact structure and length of the inserted sequence is unknown, as indicated by question marks. (c) SALK_chr2:18Mb insertion features two individual double strand breaks, around 5 kb apart. High homology between the T-DNA strands as well as the hairpin forming original DNA piece created a secondary structure (d), that was potentially excised (e) and resulted in the deletion of the ∼5kb chromosomal fragment, as shown in main text Fig. 2. Arrowheads on the red T-DNA strand show direction. The blue line represents double stranded DNA.

**Supplementary Fig. 4: ONT reads identify insertions with lower allele frequency that are not part of assembled contigs**. We applied blastn searches of all unassembled ONT reads against the utilized vector sequence, and subsequently the TAIR10 reference genome. This identified reads such as the depicted, that confirm insertion events (like SALK_059379 on Chr 4 at 10.4 Mb) not present in the assembly or BNG maps. Our blastn strategy identified chromosomal sequence (yellow), and individual alignments with the pROK plasmid sequence revealed T-strand (blue) and vector backbone (green) sequence. The pink plasmid sequence within the vector backbone shows an internal breakpoint, which was likely caused by multiple independent insertion events within the same region. Percent identity of the raw read stretch to the reference sequence are listed within the annotations.

**Supplementary Fig. 5: Variable effects of T-DNA insertions on the local chromatin environment** (a) Quantification of H3K4me3, H2A.Z and H3K27me3 at the two domains next to the T-DNA deletion in SALK_059379 is shown. (b), (c) Impact of the T-DNA integration on the local chromatin environment at *At2g36280* in SALK_059379 seedlings. Visualization (b) and quantification (c) of H3K4me3, H2A.Z and H3K27me3 occupancy around the T-DNA insertion site in Col-0 and SALK_059379 seedlings is shown as well as the schematic illustration of the gene structure of *At2g36280* (b) including the approximate localization of T-DNA insertion. (d), (e) Visualization (d) and quantification (e) of H3K4me3, H2A.Z and H3K27me3 occupancy at flanking regions of the SAIL specific WT inversion on chromosome 1. The corresponding flanking regions in Col-0 were indicated as domain 1 and 2. (f), (g), Impact of the T-DNA integration on the local chromatin environment at *At5g38850* in SAIL_232 seedlings. Visualization (f) and quantification (g) of H3K4me3, H2A.Z and H3K27me3 occupancy around the T-DNA insertion site in Col-0 and SAIL_232 seedlings is shown. Schematic illustration of the gene structure of *At5G38850* (f) indicates the approximate localization of the T-DNA insertion. (h), (i) Impact of the T-DNA integration on the local chromatin environment at *At1g32640* in SALK_061267 seedlings. Visualization (h) and quantification (i) of H3K4me3, H2A.Z and H3K27me3 occupancy around the T-DNA insertion site in Col-0 and SALK_061267 seedlings is shown as well as the schematic illustration of the gene structure of *At1g32640* (h) including the approximate localization of T-DNA insertion. (j), (k) Impact of the T-DNA integration on the local chromatin environment at *At1g23910* in SALK_061267 seedlings. Visualization (j) and quantification (k) of H3K4me3, H2A.Z and H3K27me3 occupancy around the T-DNA insertion site in Col-0 and SALK_061267 seedlings is shown. Schematic illustration of the gene structure of *At1g23910* (j) indicates the approximate localization of the T-DNA insertion. A red arrow marks the approximate location of the T-DNA integration site in the Annoj genome browser screenshots. H3K4me3, H2A.Z and H3K27me3 occupancy was profiled with ChIP-seq whereby the Col-0 IgG sample serves as a negative control. All shown AnnoJ genome browser tracks were normalized to the respective sequencing depth. The occupancy of the respective domains was calculated as the ratio between the respective ChIP-seq sample and the Col-0 IgG control.

**Supplementary Fig. 6: T-DNAs affect the local chromatin in a highly diverse manner** a), (b) Impact of the T-DNA integration on the local chromatin environment at *At5g11930* in SALK_017723 seedlings. Visualization (a) and quantification (b) of H3K4me3, H2A.Z and H3K27me3 occupancy around the T-DNA insertion site in Col-0 and SALK_017723 seedlings is shown as well as the schematic illustration of the gene structure of *At5g11930* (a) including the approximate localization of T-DNA insertion. (c) Genome browser visualizes the T-DNA-induced *de novo* trimethylation of H3K4 and incorporation of H2A.Z at in 50B_HR80 seedlings. The new T-DNA-induced domain within the gene body of *At3g57300* is indicated as well as a schematic illustration of the gene structure of *At3g57300* including the approximate localization of the corresponding T-DNA insertion. (d) Quantification of the new H3K4me3 and H2A.Z domain within the gene body of *At3g57300* is shown. (e), (f) Impact of the T-DNA integration on the local chromatin environment at *At1g01700* in SALK_117411 seedlings. Visualization (e) and quantification (f) of H3K4me3, H2A.Z and H3K27me3 occupancy around the T-DNA insertion site in Col-0 and SALK_117411 seedlings is shown as well as the schematic illustration of the gene structure of *At1g01700* (e) including the approximate localization of T-DNA insertion. (g), (h) T-DNA integration impacts the local chromatin environment at *At3g46650* in SALK_069537 seedlings. Visualization (g) and quantification (h) of H3K4me3, H2A.Z and H3K27me3 occupancy around the T-DNA insertion site in Col-0 and SALK_069537 seedlings is shown. Schematic illustration of the gene structure of *At3g46650* (g) indicates the approximate localization of the T-DNA insertion. All ChIP-seq data was visualized with the AnnoJ genome browser. A red arrow marks the approximate location of the T-DNA integration site in the Annoj genome browser screenshots. H3K4me3, H2A.Z and H3K27me3 occupancy was profiled with ChIP-seq whereby the Col-0 IgG sample serves as a negative control. All shown AnnoJ genome browser tracks were normalized to the respective sequencing depth. The occupancy of the respective domains was calculated as the ratio between the respective ChIP-seq sample and the Col-0 IgG control.

BNG: Bionano Genomics Irys optical genome mapping
CaMV 35S: cauliflower mosaic virus 35S
Col: *Arabidopsis thaliana* Columbia ecotype, wild type control
LB: T-DNA left border
NOSp: nopaline synthase promoter
NOSt: nopaline synthase terminator
ONT: Oxford Nanopore Technologies MinION long-read nanopore sequencer
PRC2: polycomb-repressive complex-2
RdDM: RNA-directed DNA methylation
RNAi: RNA interference
siRNA: small interfering RNA
sRNA: small RNA
T-DNA: transfer DNA
Ti: Tumor-inducing

